# Where’s the noise? Key features of neuronal variability and inference emerge from self-organized learning

**DOI:** 10.1101/011296

**Authors:** Christoph Hartmann, Andreea Lazar, Jochen Triesch

**Affiliations:** Frankfurt Institute for Advanced Studies, Johann Wolfgang Goethe University, Frankfurt am Main, Germany; International Max Planck Research School for Neural Circuits, Max Planck Institute for Brain Research, Frankfurt am Main, Germany; Ernst Strungmann Institute (ESI) for Neuroscience in Cooperation with Max-Planck-Society, Frankfurt am Main, Germany

## Abstract

**Abstract:** Trial-to-trial variability and spontaneous activity of cortical recordings have been suggested to reflect intrinsic noise. This view is currently challenged by mounting evidence for structure in these phenomena: Trial-to-trial variability decreases following stimulus onset and can be predicted by previous spontaneous activity. This spontaneous activity is similar in magnitude and structure to evoked activity and can predict decisions. All of the observed neuronal properties described above can be accounted for, at an abstract computational level, by the sampling-hypothesis, according to which response variability reflects stimulus uncertainty. However, a mechanistic explanation at the level of neural circuit dynamics is still missing.

In this study, we demonstrate that all of these phenomena can be accounted for by a noise-free self-organizing recurrent neural network model (SORN). It combines spike-timing dependent plasticity (STDP) and homeostatic mechanisms in a deterministic network of excitatory and inhibitory McCulloch-Pitts neurons. The network self-organizes to spatio-temporally varying input sequences.

We find that the key properties of neural variability mentioned above develop in this model as the network learns to perform sampling-like inference. Importantly, the model shows high trial-to-trial variability although it is fully deterministic. This suggests that the trial-to-trial variability in neural recordings may not reflect intrinsic noise. Rather, it may reflect a deterministic approximation of sampling-like learning and inference. The simplicity of the model suggests that these correlates of the sampling theory are canonical properties of recurrent networks that learn with a combination of STDP and homeostatic plasticity mechanisms.

**Author Summary:** Neural recordings seem very noisy. If the exact same stimulus is shown to an animal multiple times, the neural response will vary. In fact, the activity of a single neuron shows many features of a stochastic process. Furthermore, in the absence of a sensory stimulus, cortical spontaneous activity has a magnitude comparable to the activity observed during stimulus presentation. These findings have led to a widespread belief that neural activity is indeed very noisy. However, recent evidence indicates that individual neurons can operate very reliably and that the spontaneous activity in the brain is highly structured, suggesting that much of the noise may in fact be signal. One hypothesis regarding this putative signal is that it reflects a form of probabilistic inference through sampling. Here we show that the key features of neural variability can be accounted for in a completely deterministic network model through self-organization. As the network learns a model of its sensory inputs, the deterministic dynamics give rise to sampling-like inference. Our findings show that the notorious variability in neural recordings does not need to be seen as evidence for a noisy brain. Instead it may reflect sampling-like inference emerging from a self-organized learning process.

## Introduction

At first sight, the brain seems to be a very unreliable machine: neural recordings show high trial-to-trial variability on a variety of scales [1] and this variability can affect behaviour [2]. Additionally, in the absence of sensory input or motor output, neural activity is seemingly random [3].

On a closer look, however, the trial-to-trial variability decreases at stimulus onset [4] and the remaining unexplained variability can be predicted by its preceding spontaneous activity [5]. In agreement, the neural structures that do not receive recurrent input, such as the retina, display much lower neural variability [6]. Additionally, spontaneous activity is very similar to its evoked counterpart in space, time, and magnitude [7] and it has even been proposed to outline all the possible evoked states [8]. Importantly, spontaneous activity can be shaped by learning on different time scales [9–11] and can influence our decisions [12, 13]. Finally, while the brain seems to be very noisy under frequent laboratory conditions (DC input currents and room temperature), the noise is absent [14] or reduced [15–17] in more realistic conditions, and cannot account for the full trial-to-trial variability [18].

Curiously enough, the variability does not seem to stop us from integrating known information with noisy data in a statistically optimal way in many conditions [19] (but see [20]). During the last decade different authors [19, 21–23] therefore proposed that inference and behaviour should be explained by a sampling-like procedure. In this statistical approach, the uncertainty of the system is represented by successive samples over time. More specifically, at each point in time, the instantaneous activity of the relevant brain structure represents a sample from a high-dimensional probability distribution over states of the system and the world. Certainty increases by integrating these samples over a longer time. In some situations, one or two samples may already be enough to reach sufficiently good decisions [24]. This sequential sampling can attribute functional relevance to the observed variability: The spontaneous activity then simply reflects samples from the wide prior distribution and outlines the possible evoked states. When a stimulus appears, the posterior is more constrained than the prior and by this the variability of the evoked response decreases.

This theory is very appealing on this abstract level. Yet, little is known about how it might be implemented in neural circuits. So far, some authors proposed neural network implementations of sampling-based inference (e.g. [25–27]). However, they used highly constrained designs such as feed-forward architectures, winner-take-all circuits or recurrent weights dictated by the readout and often do not account for learning. Also, these models usually make use of intrinsic noise which should be seen critically given the discussion of noise above (but see [26] for a recent counterexample). Most importantly, it is not yet clear under which conditions the key features of neural variability outlined above — the origin of the sampling hypothesis — emerge from neural circuits.

In this study, we demonstrate that all the properties of neural variability described above emerge in a noise-free recurrent neural network model without explicitly modelling sampling. The network internalizes the structure of the input and produces estimates of Bayes-optimal inference with network dynamics that resemble samples from the corresponding distribution. Finally, an analysis of this model yields predictions for new experiments.

## Results

To capture the neural phenomena outlined above, we study the behaviour of our model in two different paradigms: First, we study the effect of sequence learning on spontaneous activity. Second, we model the effect of ambiguous stimuli similar to the experiment in [13]. This setting is also used to demonstrate that probabilistic inference is in principle possible in the model. These two experiments together offer enough hooks to compare the dynamics of the model to the key features of spontaneous activity and neural variability. Before doing so, we will shortly characterize the setup of the model and its basic network dynamics:

### Network model and properties

We believe that three key features underlie most of the experimental findings on neural variability: First, recurrent connectivity allows structure in spontaneous activity and an impact on evoked responses. Second, learning is essential to match spontaneous activity to the statistics of evoked activity. Finally, homeostasis is important since feedback and learning notoriously kick systems out of their healthy regimes. We capture these properties in one of the simplest models possible: The self-organizing recurrent network (SORN, [28]) is a network of interconnected excitatory and inhibitory populations of McCulloch& Pitts model neurons. The 200 excitatory neurons are recurrently connected and receive structured input sequences. Spike-timing dependent plasticity (STDP) shapes these recurrent excitatory connections. The network dynamics are kept in check by two forms of homeostatic plasticity: Synaptic normalization ensures that the summed synaptic weights onto each neuron stay the same during STDP. Intrinsic plasticity regulates the thresholds of the excitatory neurons on a slow timescale to avoid silent or overly active neurons. These networks are for example capable of capturing the statistics and fluctuations of synaptic weights during learning [29] or of learning artificial grammars with a performance comparable to humans [30].

The dynamic properties of the SORN are shown in Figure 1. The results are taken from the inference task described later in the Results. The network approaches a log-normal weight distribution, an exponential inter spike-interval (ISI) distribution and a strong correlation between the mean excitation and mean inhibition present in the network. The log-normal weight distribution is a documented property of cortical circuits [31] and the exponential ISI distribution and its coefficient of variation indicate that the individual neurons fire irregularly with Poissonian statistics, an effect often observed in neocortex [4, 32].

**Figure 1.**
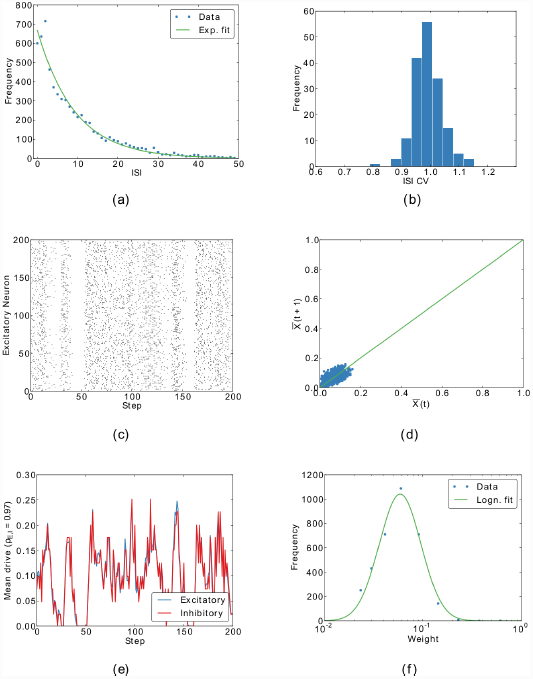
Basic properties of the network from a representative trial of the inference task. (a) The inter-spike-interval (ISI) distribution of a randomly selected neuron is well-fitted by an exponential. (b) The distribution of coefficients of variation (CVs) of the ISIs clusters around one, compatible with biological data. (c) A sample of spikes after plasticity. (d) The mean activity 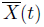 of the network is stable and does not undergo rapid changes. (e) The excitatory population receives balanced excitatory and inhibitory input. (f) After self-organization, the distribution of excitatory-to-excitatory synaptic weights (dots) approaches a log-normal distribution (solid line). (b), (c) and (d) summarize all three stimulation phases. See Fig. S1 for a similar figure for the sequence task.

**Figure 9. Figure S1.**
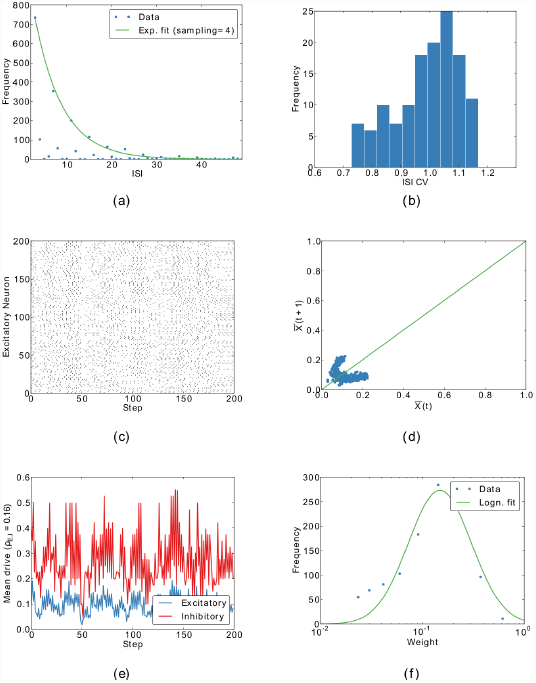
Basic properties of the network from a representative trial of the sequence learning task. (a) The inter-spike-interval (ISI) distribution of a randomly selected neuron is well-fitted by an exponential after accounting for the periodic structure of the task. (b) The distribution of coefficients of variation (CVs) of the ISIs is on average slightly smaller than for the inference task. (c) A sample of spikes from the end of the spontaneous testing phase. (d) The mean activity 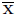 (*t*) of the network is stable and does not undergo rapid changes. (e) The excitatory population receives correlated excitatory and inhibitory input. (f) After self-organization, the distribution of excitatory-to-excitatory synaptic weights (dots) cannot be fitted by a log-normal distribution (solid line). This is due to the much longer learning time.

In the model, excitation and inhibition are balanced: For a given fraction of excitatory activity 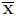, each inhibitory neuron receives on average 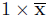 excitatory input because synaptic normalization ensures that the summed excitatory incoming weight is 1. This in turn activates the fraction 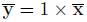 of inhibitory neurons, because the thresholds of the inhibitory units are uniformly distributed in the interval (0,1).

Taken together, the results from Fig. 1 demonstrate that this simple network readily captures some essential features of neocortex.

### Structured spontaneous activity

Spontaneous activity is structured in space and time [7, 8] and revisits those states more often that are overrepresented in natural stimuli, such as horizontal and vertical bars in [7]. This may result from adapting the spontaneous activity to the statistics of the evoked activity during development [11].

### Spontaneous activity is structured in space and time

In order to model these findings, we devised a simple experiment: During an initial self-organization phase, we stimulate the network with random alternations of two sequences: “ABCD” and “EFGH” with different relative probabilities. Presenting a letter to the networks corresponds to stimulating a corresponding subset of excitatory neurons. The letters correspond to gratings or tones in sequence learning experiments (e.g. [33]). In a second phase, we deactivate STDP to study the evoked activity of the adapted network. Finally, we stop the input and observe the network’s spontaneous activity.

Figure 2a shows the result of projecting the 200-dimensional spontaneous and evoked activity to the first three principal components of the evoked activity. These three components usually account for 40-50% of the variability. As one can see, the evoked activity captures the properties of the input by forming one activity cluster for each position in the input sequences (“A” and “E”, “B” and “F”,…). Also, the spontaneous activity closely follows the structure of the evoked activity. This observed spontaneous replay of evoked sequences is similar to [7, 9, 34]. The authors of [7] showed with optical imaging in cat area 18 that spontaneous activity is highly structured and smoothly varies over time (in their case it smoothly switches between neighbouring orientations). They also observe that spontaneous activity preferentially visits states that correspond to features that occur more often in nature (in their case horizontal and vertical bars). We demonstrate an abstract version of the later point at the end of this section.

**Figure 2.**
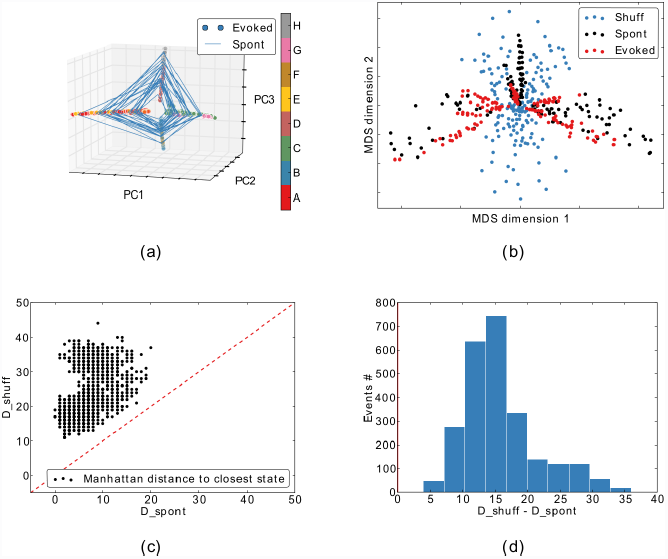
Structured spontaneous activity. The network was stimulated with the words “ABCD” (67%) and “EFGH” (33%). (a) The spontaneous activity follows the spatiotemporal trajectories of the evoked states in the PCA projection. (b) In the multidimensional scaling projection, one can see that the spontaneous activity (black) outlines the possible evoked responses (coloured). The evoked states are always closer to the spontaneous states than to the shuffled spontaneous states (c) and (d).

### Spontaneous activity outlines sensory responses

Next, we tested if the SORN model captures the finding of [8] that “spontaneous events outline the realm of possible sensory responses”. To model the conditions of the original experiment, we compared spontaneous activity to evoked activity from only 5 randomly selected letters from the original words. This captures the fact that only a subset of the “lifetime experience” of stimuli was presented during the experiment. As in [8], 150 randomly selected spontaneous events, their shuffled versions and 150 of the just described evoked events were then reduced from the high-dimensional activity patterns of excitatory neurons to 2D by multidimensional scaling (MDS). Simply put, this method uses all variability in the data to represent the distances between data points in their high-dimensional space (neural activity vectors) in the plotted two dimensions as good as possible. This is in contrast to the previous 3D-PCA plots, which only consider the variability in the first three principal components. Figure 2b shows that the spontaneous activity outlines the evoked activity while the shuffled activity cannot capture its structure. This is confirmed in Figures 2c and 2d where we show that evoked events are significantly closer to the spontaneous events than to the shuffled ones (*p* < 0.01, Wilcoxon signed-rank test).

### Spontaneous activity adapts to evoked activity

Finally, we compared the effect of learning to results from [11]. They showed that during development, the difference between the distribution of spontaneous activity and the distribution of evoked activity decreases. Interestingly, the difference decreases both for stimuli that the animal was exposed to during development (natural movies) and to artificial stimuli (gratings and bars). We try to capture the essentials of this experiment by presenting the same two sequences during the self-organization phase. After learning, we then either present the same sequences (natural condition) or the reversed sequences (control condition) for as many steps as the number of bins in the original paper. The evoked network states were then compared to the spontaneous states using the KL-divergence. As one can see in Fig. 3, our model shows a qualitatively similar behaviour in that the KL-divergence between the distribution of evoked responses in the natural condition and the distribution of spontaneous responses decreases during learning. This decrease is larger compared to the decrease observed for the reversed sequence.

**Figure 3.**
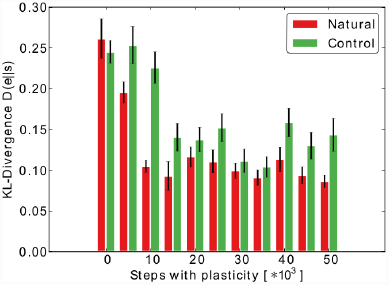
Spontaneous activity becomes more similar to evoked activity during learning. After different time periods of self-organization to “ABCD” and “EFGH”, spontaneous activity was compared to the evoked activity from the imprinted sequences (natural) or reversed sequences (control). Error bars represent SEM over 20 independent realizations.

Taken together, these results show that in our simple model the spontaneous network activity outlines the possible sensory responses after self-organization. It is important to note that none of these effects occur in a random network without plasticity (results not shown).

### Self-organization captures stimulus probabilities

Having compared our model dynamics to qualitative features of spontaneous activity, we next do a quantitative analysis on how the learnt sequences are represented in the network. In order to do so, we match the states from the spontaneous phase to the closest evoked state (see Methods for details). We assign to each spontaneous state a stimulus letter based on the stimulus of the closest matching evoked state. From this, we compute the frequency of word-occurrence in the spontaneous activity. As can be seen in Fig. 4a and 4b, the spontaneous states resemble the evoked states in two ways: First, the letters that were presented more often in the evoked activity also occur more often in the spontaneous activity. Second, the transition between states occurs in the correct temporal direction while reversed transitions rarely occur in the spontaneous activity. By varying the probability of each sequence during self-organization, we can quantify how these priors are captured by the spontaneous activity. As one can see in Fig. 4c and Fig. 4d, the probability of the words and letters are proportional to their frequency during learning. However, we observe a tendency to overrepresent the more frequent stimuli.

**Figure 4.**
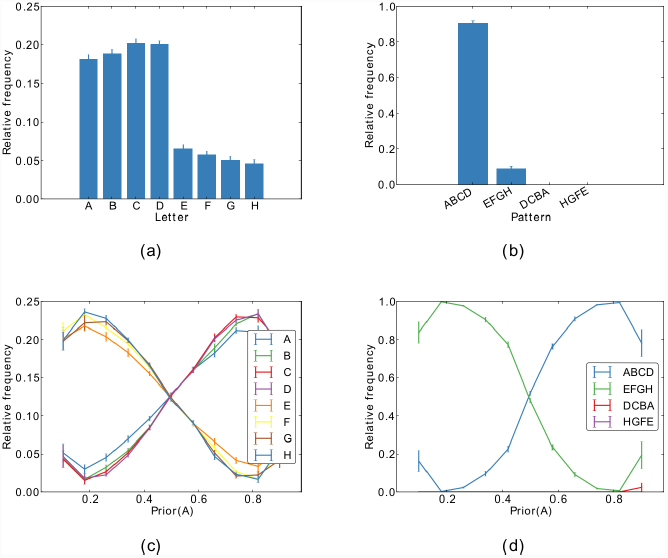
Different priors can be incorporated in the network. During self-organization, the sequences “ABCD” and “EFGH” were shown with the probabilities of Fig. 2. This is reflected in the relative occurrence of (a) each letter and (b) each word in the spontaneous activity. For different priors during self-organization, this results in the frequencies in (c) for each letter and in (d) for each word. Both show overlearning effects in that their frequencies are biased in favour of the word that was shown more often. The backwards trend with high variance at the ends can be accounted for by pathological network dynamics for some simulations with these extreme priors. Error bars represent SEM over 20 independent realizations.

### Neural variability in an inference task

After observing the properties of spontaneous activity, we next investigated the interaction between spontaneous activity and evoked activity in an inference task. Studies demonstrate that that the neural variability significantly drops at the onset of a stimulus [4], that the evoked activity can be linearly predicted from the spontaneous activity [5], and that the spontaneous activity before stimulus onset predicts the decisions when a noisy [12] or ambiguous [13] stimulus is presented.

### Modelling inference on ambiguous stimuli

We model these findings by training our networks on a task inspired by [13]. In their setting, an ambiguous face-vase stimulus is immediately followed by a mask. After the mask, the subjects have to decide whether they perceived a face or a vase.

We model this as follows: During training, the network is presented with two randomly alternating stimuli: “AXXX_ _ _…” with probability *p*_A_ and “BXXX_ _ _…” with probability 1 − *p*_A_. “A” and “B” stand for the face and vase and the common “XXX” for the mask. This is followed by a period of 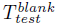 steps without stimulation, represented by the “ _”. This initial self-organization period can be seen as acquiring representations of a pure face and vase and a sense of their relative frequency of occurrence.

After self-organizing to these stimuli, STDP is switched off and a linear readout is trained to postdict whether “A” or “B” has been presented based on the neural activity at the first blank stimulus, “_”. The mask forces the network to retain an internal representation of the cue and avoids direct input-output mapping. In the test-phase, the ambiguous face-vase stimulus is modelled by stimulating the network with a mix of “A” and “B”. This is done by using 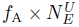 input units of “A” and 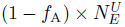 input units of “B” where *f*_A_ is a fraction. As in the original study, the delay between stimuli is random. We model this by randomly adding between 0 and 5 delay steps to the fixed delay during testing, 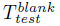, (see Methods for details).

As in the previous section, we will first investigate the qualitative behaviour of the model self-organizing to these stimuli and compare them to the experimental findings. Thereafter, we will demonstrate that the network can approximate optimal inference for the ambiguous stimuli.

### Stimulus onset quenches variability

We utilize the trial structure of this model to compare it to many features of neural variability. First, we replicate in Fig. 5 the “widespread cortical phenomenon” that the “stimulus onset quenches variability” [4]: While the neural activity and the activity in this model is in general highly variable and shows signatures of a Poisson process (cp. Fig. 1), for the same conditions there is a drop in variability measured by the Fano factor (FF) in response to stimulus onset (cp. Fig. 5). We also found that this effect is significantly stronger when the respective stimulus had a higher presentation probability (Fig. 5c vs. Fig. 5d).

**Figure 5.**
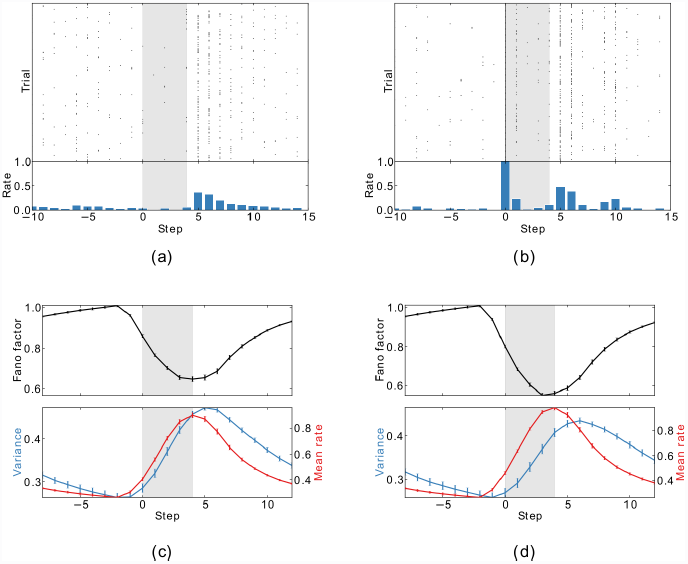
Stimulus onset quenches variability. (a) and (b) Sample spike trains from two randomly chosen neurons from the simulation in Fig. 1 aligned to the stimulus presentation (shaded area). (c) and (d) The population average of the Fano factor (FF) decreases with stimulus onset. The FF is only computed for units that do not receive direct sensory input. These results mimic Fig. 5 of [4]. FFs, mean rates and variances are for moving windows of 5 time steps. Therefore, they change before stimulus onset. (c) was computed for the presentation of stimulus “AXXX_ _ _..” during the test phase given that it had a probability of 0.1 during self-organization. (d) In turn, “BXXX_ _ _…” had a probability of 0.9 in the same experiment. Error bars represent SEM over 20 independent realizations.

As one can see in the bottom plots of Fig. 5c, the rise of the FF is accompanied by a rise of the mean firing rate. To control for this, we performed a “mean matching” analysis as suggested in [4] (see Methods for details). Fig. S2 shows that a sharp decrease of the FF persists but that the amplitude is about half as strong.

**Figure 10. Figure S2.**
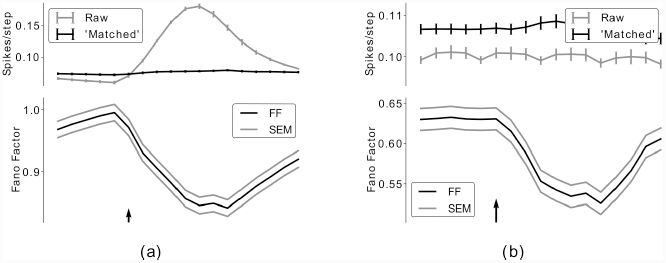
Mean-matched Fano factors. To control for effects of the mean, we computed the Fano factors with the mean-matching method proposed in [4]. This was done for both (a) the inference task and (b) the sequence learning task. As in the original paper, we averaged over all conditions. Therefore, we cannot distinguish between stimuli as we did in Fig. 5. Both conditions used the same prior as in Fig. 7 and Fig. 2. Error envelopes are SEM over 20 independent realizations.

Interestingly, this increase of the mean firing rate is higher for the stimulus that was presented more often. This matches a recent study on sequence learning with gratings in V1 [33]. It reported that when a sequence is presented for a fixed amount of trials in a passive viewing task, the response magnitude is significantly higher than when all permutations are randomly alternated during the same number of trials.

### Spontaneous activity predicts evoked activity and decisions

Next, we set out to replicate two similar findings: First, the authors of [5] found in a first groundbreaking study on the interaction of spontaneous activity with evoked activity that the optical imaging response evoked by simple bar stimuli is almost identical to the sum of the spontaneous activity prior to stimulus onset and the average stimulus-triggered response. Second, the original study that we model here [13] found that spontaneous activity prior to stimulus onset predicts the decision of the subjects. In a way, both studies show that spontaneous activity has predictive power for the evoked response.

We replicate these findings by training simple linear classifiers either to predict the evoked spiking of individual cells from the spontaneous activity immediately before stimulus presentation, or to predict the decision of the network after the presentation of stimuli with different ambiguities. This is compared to a baseline prediction that is based on shuffled spontaneous activity.

We find that the spontaneous activity prior to stimulus onset allows prediction of the evoked activity (Fig. 6a) and the final decision (Fig. 6b) in a linear manner. Please note that the baseline in Fig. 6b is not at 50%. This is because the decisions of the network are usually biased towards “A” or “B”. This can be facilitated by the control classifier using only the bias term. These two complementary experiments demonstrate that the network’s spontaneous activity prior to the stimulus contains significant information about subsequent evoked responses and behaviour as demonstrated experimentally in [5] and [13].

**Figure 6.**
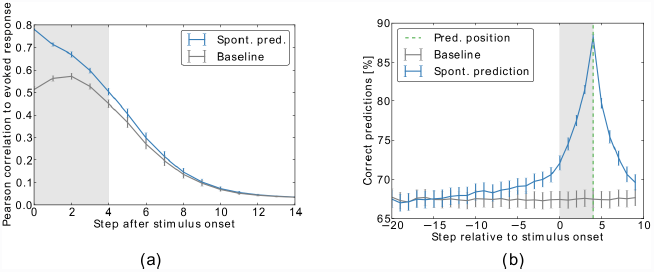
Spontaneous activity has predictive power. (a) Trial-to-trial variability is well predicted from activity prior to stimulus onset. The figure shows the correlation between the variable evoked response and either a linear prediction based on the spontaneous activity state prior to stimulus onset (blue) or on the trial-shuffled state (baseline, grey). Similar to [5], Fig. 4, the decay of the correlation starts out linearly and then proceeds to decay exponentially. The prediction was computed for the prior displayed in Fig. 7a. (b) The decisions of the network can be predicted from previous spontaneous activity. The plot shows the overlap between actual network decisions and network decisions predicted by activity surrounding the decision. The grey line corresponds to predictions from shuffled spontaneous activity. Predictions are averaged over all priors of Fig. 7b. Error bars represent SEM over 20 independent realizations.

Taken together, all these results show that the complex dynamics of a very simple self-organizing recurrent neural network suffice to reproduce key features of neural variability.

### Sampling-like inference emerges from self-organization

Our model self-organizes during repetitive presentations of two input sequences. In the current section, we investigate how the decisions of the model depend on the frequency of presentation of each sequence during self-organization (prior probability) and their ambiguity during testing. We already know from Fig. 4 that priors can be incorporated into the network. The interesting question now is how these interact with evoked activity.

Taking a Bayesian approach, we can define a simple model for combining the prior and the stimulus ambiguity, which we interpret as a likelihood here:

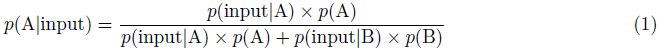

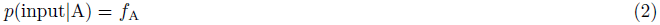

In a sampling framework, the decisions of the network for “A” should occur with frequencies similar to its posterior *p*(A|input). This is at odds with other ways to represent probabilities. For example, the posterior could be coded in the relative difference between readout activities for “A” and “B”. This would result in identical decisions for identical ambiguities. Consequently, the network would always decide for the stimulus with the higher posterior probability.

As one can see in Fig. 7a, the fraction of decisions for either “A” or “B” are an approximation of the posterior. This indicates that the network mimics sampling to represent the posterior. Fig. 7b shows that this holds for different priors. We believe that two factors interact in the network to learn the above likelihoods and the model. First, firing thresholds of the input units are regulated by intrinsic plasticity. This entails that the “A”-neurons will not fire every time “A” is presented because they might have fired too frequently in the past. Second, input units (and neurons further downstream) for stimulus “A” can be active when the network is presented with stimulus “B” due to the recurrent circuitry. These two mechanisms ensure that the network already has to deal with ambiguities during the self-organization and learning phase.

**Figure 7.**
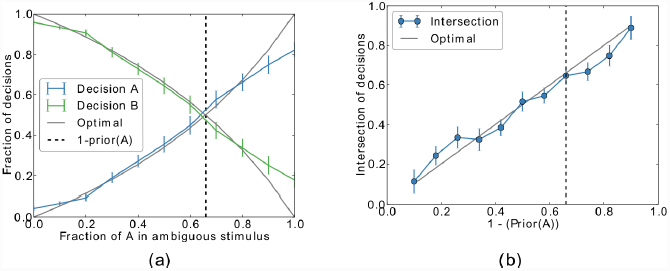
Performance of the network in the inference task. (a) The network self-organizes during repeated presentations of the sequences “AXXX_ _ _…” (33%) and “BXXX_ _ _…” (67%) with blank intervals in between. In the test phase, an ambiguous mix of cue “A” and cue “B” is presented. The fraction of “samples” (i.e. decisions for “A” or “B” at the first blank state “_”) approximates an optimal integration of cue likelihood and prior stimulus probability for the given prior (dashed line). (b) The intersections of the decisions for different priors. The prior from (a) is again represented by the dashed line. Error bars represent SEM over 20 independent realizations.

One should note, however, that longer learning will eventually drive the network towards the stimulus that was presented more often similar to the results in Fig. 4d. The intersection line in Fig. 7b will then take on a sigmoidal shape. The cause for this overlearning will be analyzed in the next section.

### Network analysis

Given all these data, the question arises how the underlying mechanisms interact to give rise to this variety of features. To better understand the dynamics of the neurons, we determine the conditional probability for neuron *x*_*i*_ to spike given that neuron *x*_*j*_ spiked at the prior time step when no input is presented. This will elucidate both how the excitatory connectivity affects the network dynamics and how the network dynamics affect the connectivity via STDP.

### Single-cell analysis

We can see in Fig. 8a that the conditional probability of neuron *x*_*i*_ spiking given that neuron *x*_*j*_ spiked at the previous time step (*p*(*x*_*i*_ = 1|*x*_*j*_ = 1)) grows roughly linearly with the synaptic weight 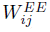 between both neurons except for saturation effects: 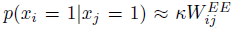. This relation, as simple as it might seem, immediately breaks down if IP or SN are deactivated (Fig. S4): While the general and intuitive trend persists that high weights imply higher conditional firing probabilities, the linear relation vanishes. The conditional probabilities in these figures were always computed for the spontaneous phase after self-organization to avoid effects from the input.

**Figure 8.**
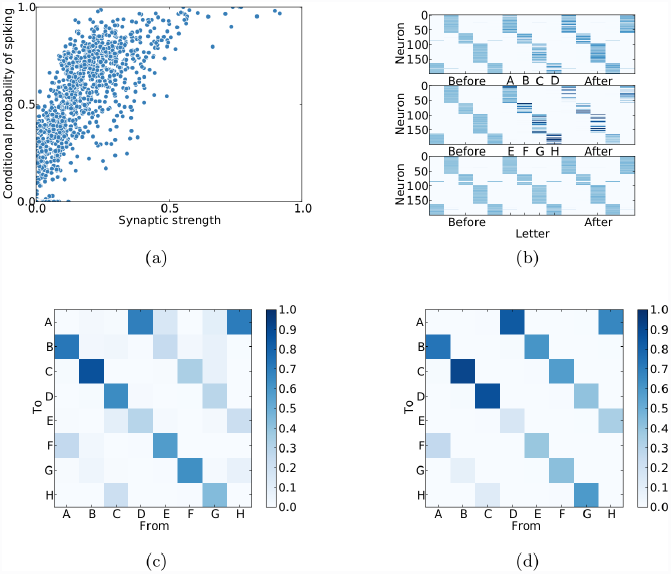
Network analysis for the sequence learning task. (a) The conditional probability of spiking *p(x*_*i*_(*t* + 1) = 1|*x*_*j*_(*t*) = 1) is roughly proportional to its synaptic weight 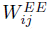 except for saturation effects for very large weights. (b) The firing probabilities of each neuron relative to stimulus onset. The network develops sequential activity patterns during the presentation of both sequences (top and middle) and during spontaneous activity (aligned to spontaneous states corresponding to “C”, bottom). Neurons were sorted according to their maximal firing probability relative to the sequence “ABCD”. (c) The prediction of transition probabilities during spontaneous activity from the singular value decomposition of **W**^*EE*^. (d) The actual transition probabilities during spontaneous activity.

**Figure 11. Figure S3.**
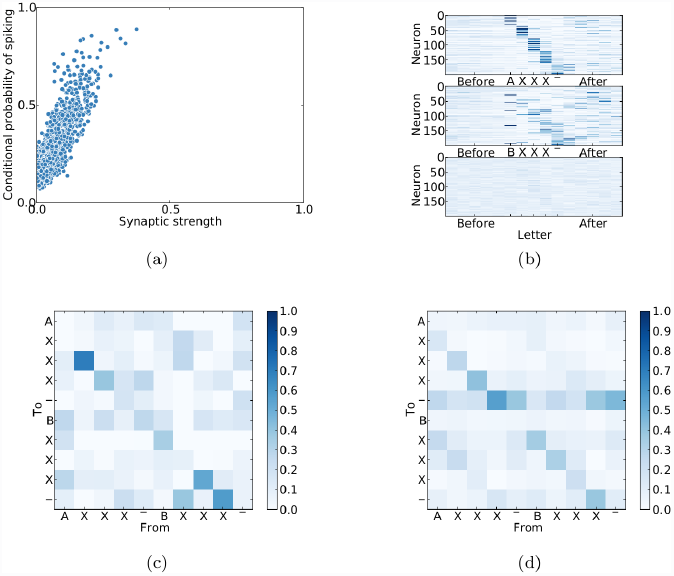
Network analysis for the inference task. (a) The conditional probability of spiking *p(x*_*i*_(*t* + 1) = 1|*x*_*j*_(*t*) = 1) is roughly proportional to its synaptic weight 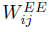. (b) The firing probabilities of each neuron relative to stimulus onset. The network develops sequential activity patterns during the presentation of both sequences (top and middle). Because the period of plasticity was not extensive as in Fig. 8, the patterns are not as similar and the spontaneous activity (aligned to spontaneous states corresponding to the second “X” of “AXXX”, bottom) is not as structured. Neurons were sorted according to their maximal firing probability relative to the sequence “AXXX_ _ _…”. (c) The prediction of transition probabilities during spontaneous activity from the singular value decomposition of **W**^*EE*^. (d) The actual transition probabilities during spontaneous activity. These plots again reflect the shorter time of STDP.

**Figure 12. Figure S4.**
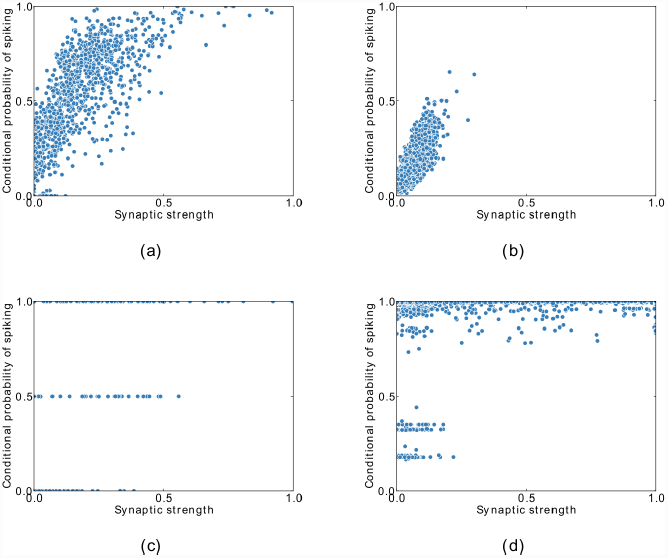
IP and SN are essential for healthy network dynamics. (a) Conditional firing probabilities from Fig. 8a. (b) The original simulation without STDP demonstrates that STDP is not necessary for the linear relation between synapse strength and firing probability. Also, STDP leads to stronger weights. (c) The original simulation without IP shows that IP is essential for maintaining correct firing probabilities. (d) The same holds true for a simulation without synaptic normalization.

This relation between connection strength and firing probability interacts with STDP: Given two bidirectional weights *W*_*ij*_ and *W*_*ji*_ with *W*_*ij*_ > *W*_*ji*_, the average weight update will be:

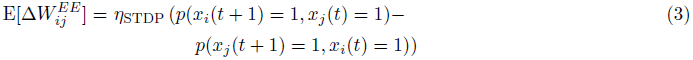

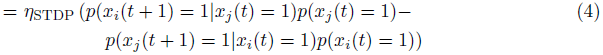

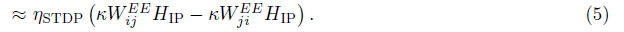

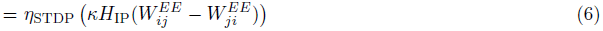

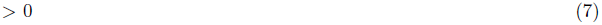

We observe that the expected weight update is directly proportional to the weight difference. In the special case of the reciprocal weight being 0, it is directly proportional to the weight. This leads to the rich-get-richer behaviour of synaptic strengths as observed in [29] and also found here in Fig. 1f. Basically, a high weight increases the firing probability of the postsynaptic neuron upon presynaptic activation, which increases the weight (Eq. 6), which increases the probability of activation, and so on. Another key factor for the rich-get-richer behaviour is synaptic normalization, as described in [29].

Apart from the skewed weight distribution, it eventually also leads to a sequential activation of neurons as observed in neocortex [35], in modelling work [36] and in the current study (Fig. 8b).

Finally, we analyse the impact of input structure on the STDP dynamics. For this, we consider the case where an excitatory neuron *x*_*A*_ receives external input with the frequency *p*(*A*). Furthermore, we assume that this neuron projects to a second excitatory input-receiving neuron, *x*_*B*_ with a very small weight, i.e. 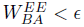, so that the impact of the weight on the firing probabilities can be neglected. This neuron receives input from the next letter in the input word, “B”. We further assume that the stimuli are presented infrequently enough so that the intrinsic plasticity does not interfere with the activation of input-receiving neurons when the corresponding input is presented, i.e. *p(A)* « *H*_*IP*_. This ensures that whenever *X*_*A*_ is activated by the input, *x*_*B*_ will be active in the subsequent time step with a probability close to 1. In this case, we have:

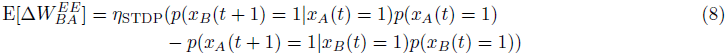

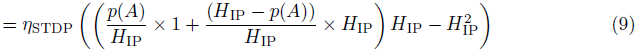

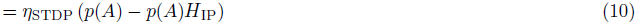

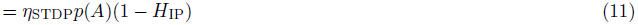

Here, we assumed independence for spiking in the reverse direction, i.e. between *x*_*B*_ (*t*) and *X*_*A*_ (*t* + 1). The first term in Eq. 9 corresponds to the conditional probability of firing when the firing is due to the stimulus 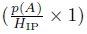 or due to the recurrent dynamics 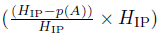.

We observe that the expected weight change is directly proportional to p(A). As a result of this, if we consider two stimuli with different frequencies, the weights will grow stronger for the more frequent stimulus. In fact, they are directly proportional so that a stimulus that is twice as frequent will have twice as strong a weight in the network. Taking into account the proportionality between the weight and the conditional firing probability, this means that the more frequently presented stimulus during learning will also have a higher probability of recall.

The above direct proportionality will break down if the weights become strong enough to have an impact on firing probabilities. This case will again result in rich-get-richer behaviour. This will lead to the overrepresentation of probabilities as observed in Fig. 4. While this specific case assumes that these neurons receive direct input, they could of course also be neurons that receive indirect input. In this case, the subsequent activation of the receiving neuron would probably not be as certain and therefore the effect less strong.

### Population analysis

Having done all these analyses on the interactions of single neurons, it is important to know if these results generalize to the population as a whole. More specifically:

1. Can STDP imprint the sequential input structure in the recurrent excitatory connectivity?
2. Do these connections correspond to the actual transition probabilities of the network states when it is running without input?

We analyse these question by simplifying the excitatory network dynamics to a linear dynamical system:

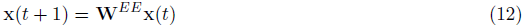

We then apply singular-value decomposition (SVD) to the recurrent weight matrix:

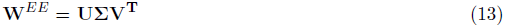

Because *U* and *V* are orthonormal and Σ diagonal, we have

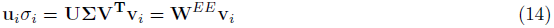

By comparing (12) and (14), one can see that the vectors **v**_***i***_ and ***u***_***i***_σ_*i*_ define transitions similar to **x**(*t*) and **x**(*t* + 1). We therefore analyse the behaviour of learned connections by matching each vector **v**_*i*_ and **u_*i*_** to their closest matching evoked state. Thereby we get from transitions between vectors to transitions between input letters. If we scale each transition by its singular value σ_*i*_ and normalize to 1, we can predict the transition probabilities.

In Fig. 8c, the result of this analysis is applied to the sequence learning task. When compared to the actual transitions during spontaneous activity in Fig. 8d, one can see that the SVD-analysis approximates the actual transition probabilities. This entails that STDP can indeed imprint the input structure in the weight matrix and that these probabilities are correctly represented during spontaneous activity. The corresponding results for the inference tasks can be found in Fig. S3.

Taken together, these analyses explain why and how the network activity acquires the stimulation structure in Fig. 2 and how the input priors and can on the one hand be imprinted into the network connectivity (cp. Fig. 4) and on the other hand utilized during testing (cp. Fig. 7).

## Discussion

A variety of phenomena connect variability of neural responses and spontaneous activity in the neocortex. However, there is growing evidence that the brain seems to be less stochastic than previously believed. We proposed that the observed variability is not simply noise but might in fact carry information and could be interpreted as samples from the Bayesian posterior.

We have shown that key properties of neural variability emerge in a simple deterministic network of recurrently connected spiking neurons that self-organize to different input patterns. We provided a proof of concept that the model is able to perform inference similar to the sampling-theoretic ideas while only making use of the internal dynamics and variability in the input presentation. Our data demonstrate that the stimulus dependence of neural variability and the structure as well as predictive power of spontaneous activity can all be seen as emergent consequences from the self-organizing dynamics of recurrent neural networks with STDP and homeostatic mechanisms. To complement these data, our analysis demonstrates that the homeostasis is essential to both learn and recall the input probabilities. These results in combination with the simplicity of our model suggest that the combination of STDP and homeostasis in recurrent networks suffices to account for the key findings on neural variability.

To explain a variety of experiments on very different scales, we decided to use a generic model. Despite its simplicity, this allowed us to reproduce data obtained from multi-electrode recordings to optical imaging and even fMRI. This implies that some of these results should be seen as a reproduction on an abstract and qualitative level: For example, we can show that the spontaneous activity obtained here is structured and has predictive power, but we did not attempt to reproduce the exact time course of the activity in the fusiform-face-area.

Also, while there is evidence for the basic plasticity mechanisms we use here in the neocortex and the hippocampus (see Methods), the unidirectional connections resulting from our STDP rule seem to be at odds with data on above-chance bidirectional connections (e.g. [37], but see [38]).

On the other hand, the generality of our model allows us to predict that the phenomena that so far have only been reported for imaging data (e.g [5, 7, 13]) are also present on the spiking level. We predict that the spiking activity prior to stimulus onset could be useful to predict evoked activity and decisions. We also predict that spontaneous spike patterns are shaped by learning and reflect the presentation probabilities of their corresponding stimuli. Another concrete prediction is that the Fano factor decreases more strongly at stimulus onset for stimuli that have a higher probability of occurring (cp. Fig. 5).

Finally, a recurring theme in this work is the overlearning found in Fig. 4 and Fig. 7 and analyzed at the end of the Results. Due to the influence of recurrent reactivation of already learnt sequences on the learning process, very frequent stimuli will tend to have a reinforcing influence during learning and thereby become overrepresented in the network. This in turn suppresses infrequent stimuli. This simple interaction seems to be an inevitable feature of learning in recurrent networks. We therefore predict that sequence learning *in vivo* (as e.g. in [33]) also does not stop at the exact relative probability but overrepresents very frequent stimuli.

While the results presented here are in themselves very encouraging, it is important to mention that previous studies demonstrated the utility of the SORN model for both computational performance and explaining biological data: [28] first introduced this self-organizing reservoir and showed that it is superior to classical, static, reservoir computing approaches when learning to predict structured input sequences. This is due to an expanded input representation and a learning of the input structure with the same combination of plasticity rules used in this work. More recently, the authors of [29] showed that a very similar network reproduces key biological data on synaptic weight statistics and fluctuations. This model accounted for both the log-normal distribution of excitatory-excitatory weights (cp. Fig. 1f) and the fluctuations of individual weights over time. Finally, an independent group recently validated this model on a grammar-learning task [30].

Recently, there is evidence from neuroscience that cortical dynamics resemble a self-organizing reservoir [39]. For example, it was demonstrated in [40] that the fading memory property present in this approach and at the basis of the field of reservoir computing can also be found in the visual cortex.

In summary, this suggests that the results found here are not artifacts from a specifically tuned network model but should rather be seen as features of a generic model that was published before some of the findings on neural variability reproduced here were reported (e.g. [4, 8, 11]).

While we are not the first to model the effects considered here, we are, to the best of our knowledge, the first to account for all these effects in unison and in such a simple model. For example, constrained balanced networks can capture the decline of the Fano factor [41, 42]. They also succeeded in capturing the effect that most recordings in [4] show Fano factors above 1, while we only observed Fano factors close to 1. One explanation for this might be the recent double-Poissonian model proposed in [32]. They showed that the additional variability can be accounted for by a second, slowly varying process like attentional states or neuromodulation. Since a process like this is missing in our model, we also do not observe this additional variability.

These balanced network models are an important contribution in showing that cellular adaptation [42] or clustering [41] might account for the characteristic decline of the FF. However, apart from using more complex models to account for this effect, the balanced networks also did not employ any learning. Thereby, they cannot to account for the other properties of neural variability treated here.

Other modelling studies tried to derive neural implementations of the sampling theory. Most prominently, the authors of [27] showed in a rigorous analysis that in a feed-forward like structure with winner-take-all circuits, STDP can be used to approximate a sampling-version of expectation maximization and thereby learn a generative model of the input. Similar work has been done in [43], who showed that spatio-temporal patterns can be entrained in a network with an artificial importance-sampling rule. A more biological approximation of this again resembles STDP. Therefore, recent studies demonstrate that STDP is a good candidate to learn appropriate weights to represent statistics of the input (see also [44]). However, all of these models make heavy use of intrinsic noise to achieve these sampling-effects (but see [26]) and do not relate their results to the biological data cited in the current work.

At this point it is important to repeat that we did not aim to model Bayes-optimal inference for arbitrary inputs. As a matter of fact, the brain, too, is not optimal in most conditions [20] and the sampling hypothesis is still highly debated (see e.g. [45] or [46]). Rather, we tested if the previously published SORN model can account for the key properties of neural variability commonly used as evidence for sampling. Interestingly, we found sampling-like inference in this network, which suggests that this is a generic property of recurrent circuits learning with STDP and homeostasis.

As just described, many models of recurrent neural networks make heavy use of intrinsic noise. This is not only practical for modelling sampling but also to stabilize networks in irregular regimes or to avoid oscillations or epilleptic-like behaviour. It is usually justified by referring to the data on neural variability in cortex. We demonstrated here that the data on neural variability can be accounted for by deterministic intrinsic dynamics and a combination of plasticity mechanisms. Due to these plasticity mechanisms the network still functions in irregular regimes and is able to approximate probabilistic inference. This suggests that the neocortex does not have to be that noisy after all. In fact, many biological studies suggest that action potential generation is highly deterministic and synaptic transmission becomes very reliable as long as experiments are performed under appropriate temperatures and the sum of synapses from one neuron on the other are taken into account [14–16]. Therefore, we propose that the common practice to make heavy use of noise in neural simulations should not be taken as the gold standard and alternative dynamical approaches should be more thoroughly investigated.

Still, observing probabilistic inference in a deterministic network seems self-contradictory. However, recent work showed that Boltzmann Machines can be approximated by a sufficiently chaotic system [47]. Such chaos can emerge from deterministic neural networks, for example by balanced excitation and inhibition [48].

We hypothesize that a similar regime might emerge in our work from the intrinsic plasticity rule. This rule always slightly shifts the individual thresholds of each neuron and thereby might facilitate dynamics with sufficient fluctuations [49]. Simply put, without intrinsic plasticity the neurons would fire with a constant rate derived from the network connectivity and their thresholds. However, the intrinsic plasticity rule forces the neurons to fire with a different mean rate. Thereby, they deviate from the regular activity patterns and introduce perturbations in the network, which could eventually make the network irregular. Finally, because the target rates are different for each neuron, synchronous activity is not possible for a long time which might lead to asynchronous irregular firing. Of course, the intrinsic plasticity rule does not have a direct counterpart in the cortex, but many mechanisms such as refractory periods, spike rate adaptation, and “real” intrinsic plasticity [50] have effects very similar to the ones just described.

We are currently studying the exact influence of intrinsic plasticity and its impact on sampling-like inference and therefore leave this for future work.

Also, we do not claim that the brain is entirely noise free. But our results show that a completely deterministic model performing self-organized learning and inference is *sufficient* to account for the key findings about neural variability. Adding small amounts of noise to these networks does not seems to change their behavior drastically, but more research is needed to study the effects of different kinds of noise to these networks more thoroughly.

All in all, we have shown that trial-to-trial variability in cortical recordings does not necessarily originate from intrinsic noise. Rather, the unexplained variability might reflect a deterministic approximation of sampling-like learning and inference. This entails that the “noise” actually contains valuable information about the current context which is exploited to make inferences about the world.

## Materials and Methods

Our model is based on the self-organizing recurrent network (SORN) [28]. Pilot studies related to the work presented here have been presented at a conference [51]. For more detailed information and a validation of the results, the code will be made available on the website of the authors^1^. The exact parameters described in the text are listed in Table 1.

**Table 1.** Parameters used in the sequence learning task and in the inference task.

| Parameter | Sequence | Inference |
| --- | --- | --- |
| $N^E$ | 200 | 200 |
| $N^U$ | 10 | 10 |
| $p^{EE}$ | 0.1 | 0.1 |
| $\eta_{\text{STDP}}$ | 0.001 | 0.001 |
| $\eta_{\text{IP}}$ | 0.001 | 0.001 |
| $H_{\text{IP}}$ | 0.1 | 0.1 |
| $\epsilon_{\text{IP}}$ | 0.01 | 0.01 |
| $T_{max}^i$ | 0.35 | 1.0 |
| $w^{in}$ | 0.5 | 0.5 |
| $T_{\text{plastic}}$ | 20000 | 1500 |
| $T_{\text{train}}$ | 20000 | 20000 |
| $T_{\text{test}}$ | 50000 | 50000 |
| $T_{\text{plastic}}^{\text{blank}}$ | 0 | 5 |
| $T_{\text{train}}^{\text{blank}}$ | 0 | 5 |
| $T_{\text{test}}^{\text{blank}}$ | - | 15 |

### Network Model

The network consists of a population of *N*^*E*^ excitatory and *N*^*I*^ = 0.2 × *N^E^* inhibitory McCulloch-Pitts threshold neurons [52]. The connections between the neurons are described by weight matrices where, for example, 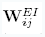 is the connection from the *j*^th^ inhibitory neuron to the *i*^th^ excitatory neuron. We model the excitatory to excitatory connections as a sparse matrix with a directed connection probability of *p*^*EE*^ and no excitatory autapses 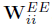. **W**^*EI*^ as well as **W**^*IE*^ are dense connection matrices. We do not model connections within the inhibitory population. All weights are randomly drawn from the interval [0,1] and then normalized by the synaptic normalization described below.

The input to the network is modelled as a series of binary vectors **u**(*t*) where at each time step *t* during input presentation all units are zero except for one. For better readability we assign an arbitrary letter to each such state when describing different input sequences later on. The letter “_” corresponds to no input presentation. Each input unit *u*_*i*_ projects to *N*^*U*^ excitatory neurons with the constant weight *w*^*in*^. These randomly selected and possibly overlapping projections are stored in *W*^*EU*^.

At each discrete time step *t*, these variables contribute to the binary excitatory state x(*t*) *∈* {0,1}^*N*^*E*^^ and inhibitory state y(*t*) *∈* {0,1}^*N*^*I*^^ as follows:

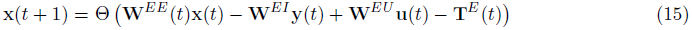

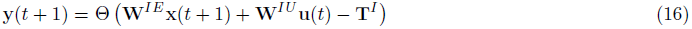

Here, Θ(a) is the element-wise heaviside step function, which maps activations **a** to binary spikes if the activation is positive. **T**^*E*^ (*t*) and **T**^*I*^ stand for the individual thresholds of all neurons and are crucial in regulating the spiking. They are randomly initialized in the interval (0,0.5) for the excitatory thresholds and 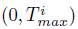 for the inhibitory ones. 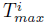 directly influences the amount of inhibition. A high maximal threshold will lead to less neurons with a low threshold and therefore less inhibition for the same amount activation. This parameter is tuned to avoid oscillations that can occur with too little inhibition but at the same time to allow computation which can be hindered by too much inhibition. The excitatory thresholds as well as **W**^*EE*^ (*t*) are adapted over time by three basic plasticity mechanisms.

It is important to note that at no point any noise is added to the network. The only external variability is the random alternation of input words, as described in the results.

### Plasticity Mechanisms

The network employs two different kinds of plasticity: Spike-timing dependent plasticity (STDP) [53–55] extracts structure from the input by shaping the weights within the excitatory population. This is counterbalanced by two forms of homeostatic plasticity: intrinsic plasticity (IP) [56, 57] and synaptic normalization (SN) [50, 58, 59]. All these mechanisms are known to co-occur in the hippocampus and neocortex.

STDP strengthens the connection 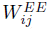 from unit *j* to unit *i* whenever a spike in *i* follows a spike in *j* (i.e. *j* helped to cause *i*) and is weakened whenever a spike in i precedes a spike in *j*. This results in the following update equation:

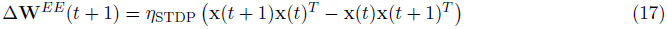

The authors of [59] showed in an EM study that the summed synaptic area per *μm* of dendrite is similar before and after LTP while the synaptic area per synapse increases and the number of synapses per *μm* decreases. This indicates that plasticity redistributes weights to avoid uncontrolled growth. This is modelled by normalizing all incoming connections of a neuron to 1 — a process called synaptic normalization (SN):

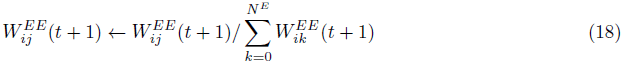

At the same time, there are a variety of regulatory mechanisms that control neural firing at different time scales such as absolute and relative refractory periods, spike rate adaptation and intrinsic plasticity [50, 57]. Here, we abstract from those by using a simple homeostatic regulation of the spike threshold at a single time scale. This intrinsic plasticity (IP) rule regulates the individual thresholds so that on average, the excitatory neurons will fire according to their target rates **H**_IP_. These target rates are uniformly drawn from the interval (***H***_IP_ − ε_IP_, *H*_IP_ + ε_IP_) for stability reasons.

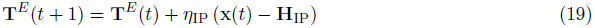

The inhibitory connections are scaled during the initialization phase so that the sum of excitatory weights received by each inhibitory unit and the sum of inhibitory weights received by each excitatory unit is 1.

The STDP and IP rules are only operating on the excitatory neurons for two reasons. First, by restricting plasticity to only the excitatory population, the model becomes simpler and thereby easier to understand and interpret. Second, there is much less data about plasticity in inhibitory neurons.

### Stimulation Paradigm

The stimulation paradigm is very similar across all tasks and can be divided into three phases:

In the self-organization phase, the network is stimulated for *T*_plastic_ steps. During this time, all plasticity mechanisms are active and the network can self-organize in the presence of given input. As stated above, each allowed input vector will from now on be represented by a letter. The stimulation paradigm can then for example be a random alternation of the words “ABCD” and “EFGH”. After this, STDP is switched off so that the properties of the learnt connections can be studied without interference from continued changes to the weights. This is done during a training and testing phase.

In the training phase, the stimuli are kept identical to observe the properties of the network under the self-organization conditions and training appropriate readouts for *T*_train_ steps.

Then, during the testing phase, we either observe the spontaneous activity while the network is not stimulated, or we test our readout on data generated by stimulating the network with appropriate stimuli for *T*_test_ steps.

For these phases, the intrinsic plasticity is still active to ensure stable average activity. After each phase, the network state is randomly reset in accordance with [49]. To simulate the trial-like structure of typical experiments, we chose to have blank periods between each stimulus presentation. The length of these periods are 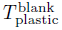 and 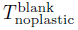 for the phases with and without plasticity.

### Analysis Methods

In this study, we compare the behaviour of our model to a variety of experimentally reported results. While our network is operating on a timescale of roughly 25ms (the width of a typical STDP-window) and on the spiking level with rates around 4Hz, the experimental data are fMRI-BOLD signals, optical imaging data and multi-unit activity spike trains and have effects on time scales that range between milliseconds and seconds. Comparing these partly very different data can therefore sometimes be only done on an abstract and qualitative level. In the following section, we outline how each comparison was performed.

### Principal component analysis and multidimensional scaling

For the principal component analysis (PCA) of the spontaneous and evoked states, we did a PCA of the last 2500 steps of the training phase. We then projected these states in the subspace defined by the first three components of the PCA and also projected the same amount of spontaneous activity states in the same subspace. The spontaneous states were taken from the end of the testing phase to avoid artifacts due to the re-adaptation of the thresholds after cutting the input.

Multidimensional scaling (MDS) was performed as closely as possible to the method used in [8]. As in that paper, we used 150 points of spontaneous, shuffled spontaneous and evoked samples. These samples were chosen randomly from the end of the training and testing phase. We did not subsample our neurons to the 45 that they recorded from because the distances between individual activity patterns become too similar in that case. This did not happen for their study because they used rates instead of spikes. To account for the fact that [8] only showed a subset of all the stimuli that the network was exposed to during its development we also only used a randomly chosen subset (5 letters) of the stimuli that we presented in the self-organization phase. We used the same Matlab function for MDS as the one used in the original paper (with Kruskal’s normalized stress1 criterion).

### KL-divergence of spontaneous and evoked activity

To replicate the results of [11], we recorded spontaneous and evoked activity for the same number of time steps (750.000) and randomly subsampled 16 units from our network. Please note that due to an exponential increase in of possible patterns with the number of units, more units would have soon become infeasible to analyse with conventional methods. To compute the KL-divergence we first have to estimate the probabilitiy *p(x)* for each pattern *x* from the set of the 2^16^ possible patterns *X.* In order to do so, we created a bin for each pattern and simply counted the occurrence of each pattern. Additionally, we started with a non-informative prior by assuming that each pattern *x* ∈ *X* was already observed once. This initial prior is necessary since KL-divergence is only defined for non-zero probabilities. After normalizing, we then computed the KL-divergence between evoked and spontaneous activity according to:

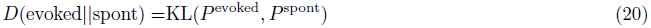

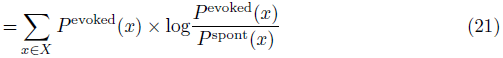

### Pattern analysis

For this analysis, we want a connection from spontaneous activity to the stimuli of the evoked activity. Given a sequence of states **x**(*t*),…, **x**(*t* + *T*) during a blank period, one would like to know how similar this sequence is to evoked patterns of activity. For this, we assigned to each spontaneous state the letter corresponding to the best-matching evoked activity state. If, for example, **x**(*t*) had the smallest Hamming distance to an evoked state when the letter A was shown, then **x**(*t*) was labelled as corresponding to input letter A. To avoid biases, the collection of evoked states used for the comparison was always obtained from the end of the training phase (the last 2500 steps). The data was further reduced by ignoring blank periods and ensuring that each letter was represented equally often in the data set.

### Inference analysis

To test whether the network can perform inference, we trained output units based on the randomly alternating presentation of the stimuli “AXXX_ _ _…” and “BXXX_ _ _…”, where “A”, “B”, and “X” each refer to a subpopulation of excitatory neurons that are stimulated whenever the letter is presented. At the presentation of “_”, no neurons receive external inputs. These stimuli model a decision task where subjects where presented with the ambiguous face-vase stimulus and had to decide whether they perceived a face or a vase after a mask [13]. “A” and “B” represent the face or vase, the following “X”s are identical for both stimuli and thereby act as a mask and delay.

To read out the model decisions after the mask, we trained a linear readout in a supervised way with least-squares regression. We performed two regressions from all neurons at the step when “_” is presented to predict either “A” or “B” (based on the recurrently maintained information of “A” or “B” that should still be present in the network). In the test phase, when ambiguous mixtures of “A” and “B” are presented, the network decision is set to “A” when the readout for “A” is larger than for “B” and the other way round. The regression target was set to 0 for all other letters from the training data, since these should not correspond to a “sampling” of “A” or “B”. The regression was performed on *T_train_* steps of evoked activity while STDP is turned off. To avoid any biases, we took equal amounts of activity samples for each letter from the end of training.

### Fano factor analysis

The Fano factor is defined as 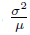. When applied to spike trains, the windowed variance *σ*^*2*^ and mean *μ* are taken over trials for each neuron, condition and time step. It is then interpreted as the variability of the data. For a Poisson process, which describes the irregular spiking of neurons in many cases quite well, the Fano factor is 1 due to the variance and mean being equal for this process.

We tried to keep our analysis of the Fano factor close to [4]. As in the original paper, the FFs were obtained by weighted regression between the spiking mean and variance over trials. To compute these two, we used a sliding window with a width of 5 bins. Together with the IP parameters, this entails a rate of 0.5 spikes per bin on average. This is comparable to the conditions of the original paper: most experiments have a rate on the order of 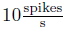 and a bin size of 50ms. One should note that due to the weighted regression and the averaging over many neurons, the resulting Fano Factor does not have to be identical to the ratio of the mean variance and the mean firing rate.

To control for an effect of the mean firing rate on the Fano factor, we also performed the “mean matching” analysis from [4]. We found that this method discards around two thirds of the data, comparable to the original study. Therefore, we only see it as a control and focussed on the real FF for the discussion.

### Prediction of evoked activity and decisions

For our last comparisons, we aimed to show that the spontaneous activity in this network can be used both to predict the following evoked activity [5] and to predict the decision of the network as for example in [12, 13].

The former was done in a similar way as the inference: For each combination of stimulus condition c and time step Δ*t*_*a*_ after stimulus onset *t*_on_, we trained a linear readout that maps **x**^*c*^(*t*_on_ − 1) to **x**^*c*^(*t*_on_ − 1 + Δ*t*_*a*_). The match between the predicted evoked activity and the actual evoked activity **x**^*c*^(*t*_on_ − 1 + Δ*t*_*a*_) was then calculated by taking the pearson correlation between both states. We allowed a bias term in the regression by appending a constant to each **x**^*c*^(*t*_on_ − 1) vector. To asses the impact of the spontaneous activity on the prediction, we compared this performance to doing the same regression from shuffled spiking patterns over trials. By shuffling over trials, the statistics of the individual patterns persist, but their relation to the decision vanishes. Since this still includes the bias term, this control can capture average effects but has to deal with the same number of parameters as the original regression. Therefore, if the spontaneous activity prior to stimulus onset contains information about the following evoked activity, the prediction based on the spontaneous activity should be better, i.e. correlate more with the true activity, than the one based on the shuffled spike trains. Also, both predictions should overlap for very late states since the information from the spontaneous activity should be “washed out” by then.

To predict the decision of the network, we also used this regression approach. For this, we divided the *T*_test_ steps of network activity with decisions into two halves — one for training the readouts to predict the decision for A and B and one for testing its performance. For each step before stimulus onset and a given stimulus, an individual readout was trained. The prediction was then defined as the higher readout. We evaluate the quality of this prediction by comparing it to the actual decisions of the network and computing the agreement between both for the testing data.

## Footnotes

1 http://fias.uni-frankfurt.de/neuro/triesch/

## Acknowledgments

We would like to thank Bernhard Nessler for fruitful discussions.

